# Bayesian estimation of cell-type-specific gene expression per bulk sample with prior derived from single-cell data

**DOI:** 10.1101/2020.08.05.238949

**Authors:** Jiebiao Wang, Kathryn Roeder, Bernie Devlin

## Abstract

When assessed over a large number of samples, bulk RNA sequencing provides reliable data for gene expression at the tissue level. Single-cell RNA sequencing (scRNA-seq) deepens those analyses by evaluating gene expression at the cellular level. Both data types lend insights into disease etiology. With current technologies, however, scRNA-seq data are known to be noisy. Moreover, constrained by costs, scRNA-seq data are typically generated from a relatively small number of subjects, which limits their utility for some analyses, such as identification of gene expression quantitative trait loci (eQTLs). To address these issues while maintaining the unique advantages of each data type, we develop a Bayesian method (bMIND) to integrate bulk and scRNA-seq data. With a prior derived from scRNA-seq data, we propose to estimate sample-level cell-type-specific (CTS) expression from bulk expression data. The CTS expression enables large-scale sample-level downstream analyses, such as detecting CTS differentially expressed genes (DEGs) and eQTLs. Through simulations, we demonstrate that bMIND improves the accuracy of sample-level CTS expression estimates and power to discover CTS-DEGs when compared to existing methods. To further our understanding of two complex phenotypes, autism spectrum disorder and Alzheimer’s disease, we apply bMIND to gene expression data of relevant brain tissue to identify CTS-DEGs. Our results complement findings for CTS-DEGs obtained from snRNA-seq studies, replicating certain DEGs in specific cell types while nominating other novel genes in those cell types. Finally, we calculate CTS-eQTLs for eleven brain regions by analyzing GTEx V8 data, creating a new resource for biological insights.

## Introduction

Gene expression quantified at the tissue level, bulk gene expression data, has been a useful resource for understanding the etiology of different diseases. RNA sequencing technology, applied to tissue samples, is mature and its relatively cost-efficient property allows assessment of tissue from hundreds of samples, thereby producing a rich data set (GTEx Consortium, 2017; Parikshak et al., 2016; Allen et al., 2016; Wang et al., 2018; Bennett et al., 2018). However, because tissue is comprised of a variety of cell types, bulk data are the convolution of gene expression from myriad cells of various cell types. To overcome this challenge, researchers have pursued single-cell RNA-sequencing (scRNA-seq) to quantify cell-type-specific (CTS) gene expression, either at the cellular or nuclear levels (Darmanis et al., 2015; Velmeshev et al., 2019; Mathys et al., 2019). While providing important insights into etiology, such data have their own limitations: cells are typically collected from a limited number of samples, thus they lack sufficient variation over samples; and the data are noisy and technically variable due to quantification of a small number of RNA molecules. This issue is especially severe for single-nucleus RNA-seq (snRNA-seq) data from frozen tissue, which is the main specimen source for brain research. Nuclear RNA accounts for only 20-50% of the RNA molecules in the whole cell, and this fraction varies across cell types (Bakken et al., 2018). Furthermore, studies of brain tissue have found that snRNA-seq fails to detect a fraction of the microglia population (Mathys et al., 2019) and microglial activation in the human brain (Thrupp et al., 2020), yet microglia are thought to be a key cell type related to critical diseases, such as Alzheimer’s disease.

To overcome the drawbacks of bulk and scRNA-seq/snRNA-seq data, while maintaining their unique advantages, we propose to integrate bulk and single-cell data to estimate CTS expression for a large sample size. Existing methods typically can only estimate population-average CTS expression (e.g., csSAM (Shen-Orr et al., 2010)). To enable subject-level estimation, we previously developed a novel MIND algorithm (Wang et al., 2020) that extends population-average estimates to the level of subject-specific CTS by borrowing information across multiple measures of bulk level expression from the same subjects. Multi-measure bulk expression data are not commonly available, however, most datasets only have one or two measures of bulk expression per subject. Correspondingly, there have been methods developed in parallel for single-measure bulk DNA methylation data (e.g., TCA (Rahmani et al., 2019)) and gene expression data (e.g., CIBERSORTx (Newman et al., 2019)). TCA is a frequentist method similar to MIND, and CIBERSORTx relies on non-negative least squares to estimate sample-level CTS expression with the goal of separating two groups of samples. There are also cell-type analytical methods for testing the interaction of cell type fractions and the variable of interest without explicit estimation of CTS expression, such as CellDMC (Zheng et al., 2018), which was originally designed for DNA methylation data. Nonetheless, these methods have not efficiently utilized the rich information available from single-cell data.

To address these deficiencies, we develop a Bayesian MIND (bMIND) algorithm to refine the estimation of CTS expression for each bulk sample. To provide accurate and reliable estimates in this setting, we propose to use information from scRNA-seq data fully by incorporating it as prior information. Bayesian approaches are known to work well and robustly for small sample size, while incorporating prior information to regularize statistically challenging estimation. In our previous work, subject-level CTS estimates are shown to be useful for various downstream analyses, such as uncovering CTS co-expression networks and CTS expression quantitative trait loci (eQTLs) (Wang et al., 2020). Here, we will demonstrate the usefulness of bMIND by CTS differential expression (DE) analysis, assessed through simulations and by analysis of data relevant for autism spectrum disorder (ASD) and Alzheimer’s disease (AD). By comparing bMIND to other state-of-the-art methods, we demonstrate bMIND provides more accurate sample-level CTS estimates as compared to TCA (Rahmani et al., 2019) and CIBERSORTx (Newman et al., 2019) and improves power for CTS-DE analysis as compared to csSAM (Shen-Orr et al., 2010) and CellDMC (Zheng et al., 2018). Finally, with updated GTEx V8 brain data and the new bMIND algorithm, we calculate CTS-eQTLs for each of eleven brain regions and thereby create a new resource for uncovering the etiologies of complex diseases and other phenotypes.

## Results

### Bayesian estimation of sample-level CTS gene expression

To improve the estimation of sample-level CTS expression, we propose a fully Bayesian algorithm (bMIND) to incorporate prior information from single-cell data (Fig. 1). We model bulk expression of sample *i* in gene *j*, ***x***_*ij*_, for *T* ≥ 1 measures, as a product of cell type fraction (***W***_*i*_, *T* × *K*) and CTS expression (***a***_*ij*_) in Bayesian mixed-effects models

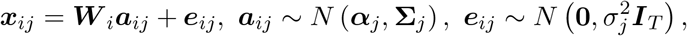

**Figure 1:**
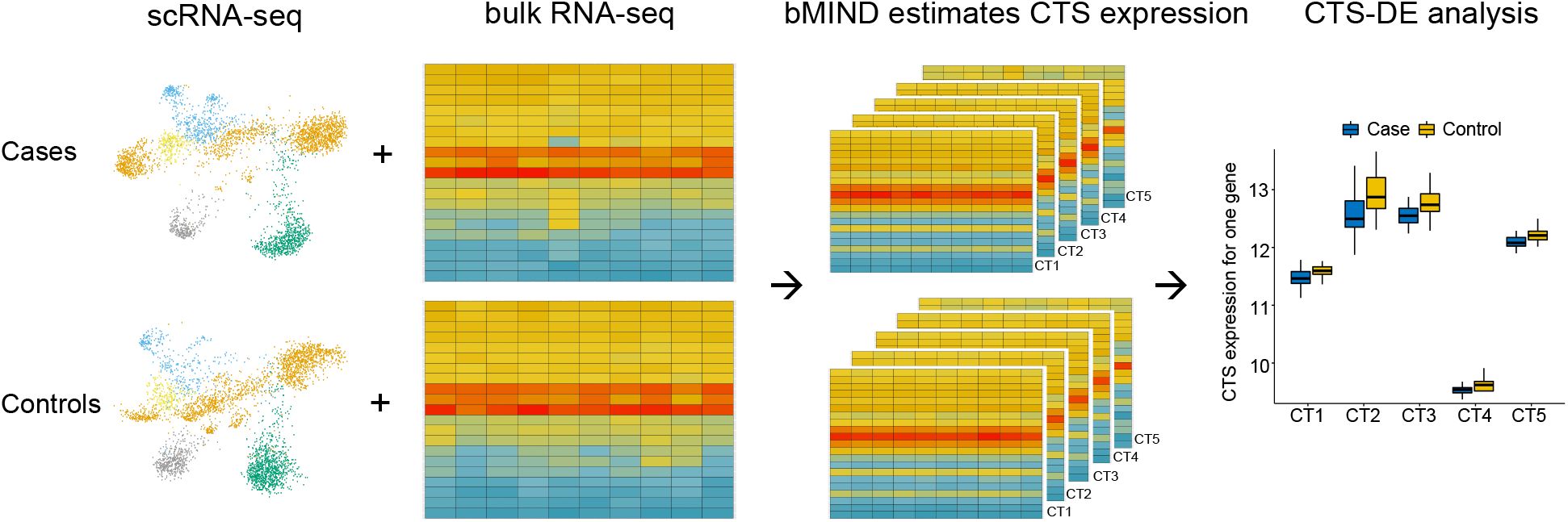
Overview of bMIND algorithm and CTS differential expression analysis (CTS-DE). With prior information from scRNA/snRNA-seq data for case and control subjects, bMIND analyzes bulk RNA-seq data and estimates sample-level CTS expression with a Bayesian approach. Here we present an example of five cell types (CT1-CT5). In the downstream analysis, as an example, we test the association between CTS expression and phenotype for each gene in each cell type and identify CTS differentially expressed genes (CTS-DEGs).

where ***α***_*j*_ (*K* × 1) is the expected CTS expression for the *j*th gene that constitutes the profile matrix, **Σ**_*j*_ (*K × K*) is the covariance matrix of CTS expression for *K* cell types, and ***e***_*ij*_ is the error term that captures the unexplained random noise with variance 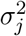. The cell type fraction (***W***_*i*_) is assumed known or pre-estimated using a cell type fraction estimation algorithm (Wang et al., 2019; Jew et al., 2020). The goal of bMIND is to provide the posterior mean of the CTS expression (***a***_*ij*_, *K* × 1) for each sample *i*, gene *j*, and *K* cell types.

To incorporate information from scRNA-seq data, we utilize these summary statistics: for each gene *j*, let 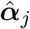 be the profile matrix, and let 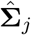 be the cell-type covariance matrix. We assume the following prior distribution.

- 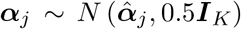, where 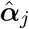 is the average CTS expression calculated from scRNA-seq data;
- **Σ**_*j*_ ∼ Inv Wishart (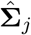, 50, where the first parameter is the expected covariance matrix and the second parameter represents the degree of belief. The inverse-Wishart distribution is the conjugate prior for the covariance matrix, which eases estimation, and it facilitates explicitly incorporating the prior covariance matrix 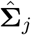 for the *j*th gene estimated from scRNA-seq data.
- 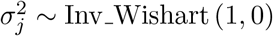, which is non-informative.

Given the technical noise and variability of scRNA-seq data, we utilize summary statistics from the scRNA-seq data rather than the raw data because summary statistics are more robust and also reduce the computation burden (Zhu et al., 2018). The hyper-parameters in the prior distributions are chosen based on empirical experiments. bMIND is robust to their specification, as shown in the Results section. We allow gene-specific parameters and analyze each gene in parallel. Although implemented with Markov chain Monte Carlo (MCMC) sampling, bMIND is computationally efficient. Depending on the sample size and number of cell types, all genes in the genome can be analyzed in approximately an hour using 30 CPU cores.

For comparison, we employ four other methods: 1) TCA (Rahmani et al., 2019), a frequentist approach similar to MIND, designed for bulk DNA methylation data, but also applicable to gene expression; 2) CIBERSORTx (Newman et al., 2019), estimating CTS expression via non-negative least squares, 3) csSAM (Shen-Orr et al., 2010), designed for microarray data to estimate population-average CTS expression, and featuring a permutation-based test for CTS-DE analysis; and 4) CellDMC (Zheng et al., 2018), designed for DNA methylation data, but applicable to gene expression data, wherein it tests CTS-DEGs by regressing bulk expression on the interaction terms between phenotype and cell type fractions. We also compare bMIND, which uses a prior derived from scRNA-seq data, with bMIND rp, a variant of bMIND, which uses a rough prior based on the analyzed bulk data.

### bMIND refines estimates of sample-level CTS expression

We evaluated the properties of bMIND with real-data analyses and realistic simulation studies. First, we checked if bMIND is able to detect variation in gene expression by cell type. We tested this by looking for consistent CTS expression across different datasets, using two existing bulk RNA-seq datasets from brain samples of subjects diagnosed with ASD and samples from unaffected subjects (Parikshak et al., 2016; Velmeshev et al., 2019) and a prior derived from snRNA-seq data (Velmeshev et al., 2019). After averaging the estimated CTS expression across samples within each dataset, for each gene, we calculated the correlation between the two averages, over cell types. The median correlation across all genes is 0.61, which increases to 0.80 when the average is restricted to marker genes, defined as the 1000 most highly expressed genes per cell type (McKenzie et al., 2018) (Fig. 2a and Supplemental Fig. S1a). For comparison, the median correlation is 0.20 and 0.27 for TCA and bMIND rp across all genes, respectively, and 0.23 and 0.34 for marker genes. Finally, we performed the same comparison by comparing snRNA-seq data (Velmeshev et al., 2019) to scRNA-seq data (Darmanis et al., 2015) and obtained median correlations of 0.46 and 0.75 for all genes and marker genes, respectively. Together, these results showed that bMIND provides meaningful estimates of gene expression profiles derived from bulk datasets.

**Figure 2:**
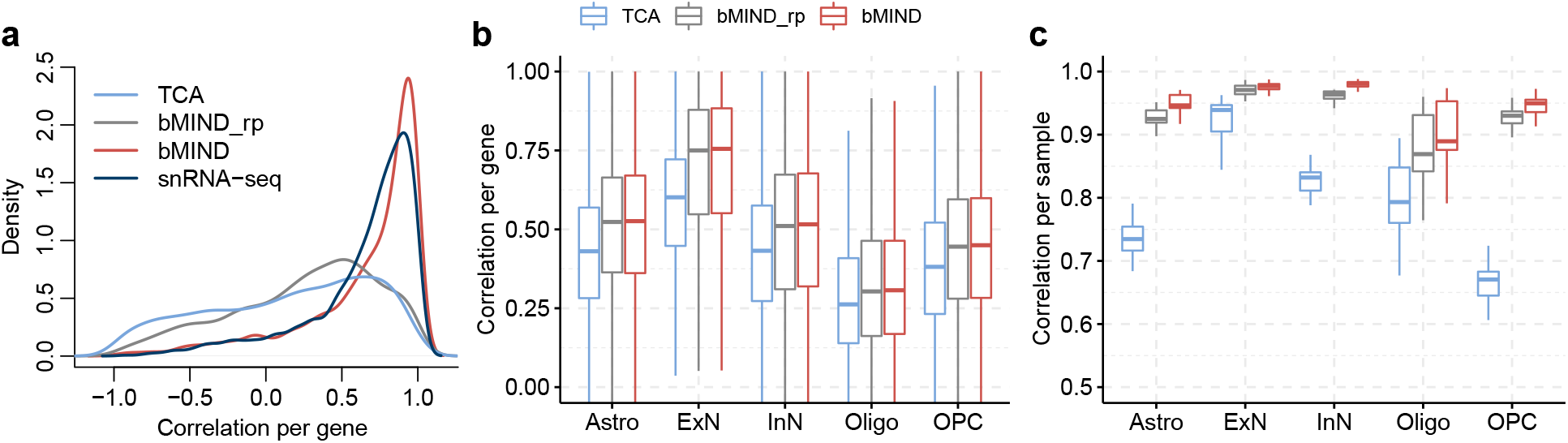
Correlation comparison for estimated sample-level CTS expression. **a**) Correlation of average CTS gene expression (across samples), over cell type, obtained from two brain RNA-seq datasets (Parikshak et al., 2016; Velmeshev et al., 2019). For a benchmark, we assess the concordance between brain snRNA-seq data (Velmeshev et al., 2019) and scRNA-seq data (Darmanis et al., 2015) (labeled as snRNA-seq). Results are provided for the top 1000 marker genes, per cell type, identified by BRETIGEA (McKenzie et al., 2018). **b,c**) Using realistic simulations, correlation between truth and estimated CTS expression are computed to compare TCA (Rahmani et al., 2019) with bMIND and bMIND rp (using a rough prior). The task involved analyzing pseudo-bulk data generated from snRNA-seq data (Velmeshev et al., 2019) obtained from ASD subjects, and bMIND uses a prior derived from the corresponding controls. For each cell type, we compute **b**) correlation across samples for each *gene*, and **c**) correlation across genes for each *sample*.

Next, we assessed whether bMIND could provide reliable sample-level CTS estimates. Ideally, this would be evaluated by comparing scRNA-seq and CTS estimates obtained from the same samples, but notably, in a comparison of bulk RNA-seq and reconstructed bulk expression obtained from snRNA-seq data, the per gene correlation was observed to be quite low (Velmeshev et al., 2019).

Presumably, due to various technical challenges and biases, snRNA-seq data do not provide a gold standard measure and this renders the desired comparison unpromising. For this reason, we used simulations to assess the correlation between estimated and true CTS expression for each cell type. Velmeshev et al. (Velmeshev et al., 2019) collected snRNA-seq data from brain tissue samples of subjects diagnosed with ASD and samples from unaffected subjects as controls. Using the single-nucleus expression for the available 41 brain samples, we grouped the nuclei into five major cell types: astrocytes (Astro), excitatory neurons (ExN), inhibitory neurons (InN), oligodendrocytes (Oligo), and oligodendrocyte precursor cells (OPC), while dropping endothelial (Endo) cells and microglia (Micro) due to low fractions. Henceforth, we shall call these the Velmeshev data. We aggregated the expression of nuclei from each ASD sample to generate pseudo-bulk data for which we know the ground truth. We estimated the prior distribution using the snRNA-seq data from control samples. After analyzing the generated pseudo-bulk data to estimate CTS expression, we calculated the correlation between the true and estimated CTS expression, per cell type, per gene. Note that by generating the bulk data from the ASD samples and prior distribution from the control samples, we assessed the robustness of the method to utilizing distinct data sources for the analysis.

The correlation per gene based on bMIND is higher than those from TCA (Fig. 2b) and CIBER-SORTx (Supplemental Fig. S1b) across cell types. Furthermore, because bMIND is a Bayesian approach, we evaluated the sensitivity of the estimates to the prior distribution specification by comparing bMIND to bMIND rp, and we found the results encouraging (Fig. 2b). Nonetheless, a precise prior can improve sample-level estimates of CTS. For instance, the correlation between estimated expression and the truth across genes, for each cell type and each sample, is considerably higher using bMIND (Fig. 2c). Finally, to show that bMIND has the ability to incorporate additional measures of bulk expression from the same samples, we conducted realistic simulations and demonstrated that more measures increase the estimation accuracy (Supplemental Fig. S1c). The bulk data were simulated with measured cell type fractions and sample-level CTS expression derived from snRNA-seq data (Velmeshev et al., 2019).

### bMIND improves power in CTS differential analysis

Having estimated sample-level CTS expression, it is straightforward to perform CTS-DE analysis (see the specific testing procedure in Methods). To evaluate performance, we conducted extensive simulation studies, assessing the false discovery rate (FDR) and power. We compared bMIND with existing methods, specifically csSAM (Shen-Orr et al., 2010) and CellDMC (Zheng et al., 2018). Other options for CTS-DE analysis are TCA (Rahmani et al., 2019) and CIBERSORTx (Newman et al., 2019); however, as reported in the literature (Jing et al., 2019), we observed inflated FDR using TCA (Supplemental Fig. S1d) and thus did not consider it in CTS-DE analysis. CIBER-SORTx (Newman et al., 2019) is not open source and thus not suitable for extensive simulation studies.

We assessed the FDR and power as a function of effect size, the number of cell types, and sample size. Under most simulation scenarios, similar to csSAM and CellDMC, bMIND controls FDR (Figs. 3a-c) at the nominal level of 0.05. More importantly, bMIND has improved power as compared to csSAM (Shen-Orr et al., 2010) and CellDMC (Zheng et al., 2018) (Figs. 3d-f). This suggests the benefits of explicitly estimating sample-level CTS expression with a Bayesian approach and flexibly accounting for FDR in the downstream sample-level CTS-DE analysis. The advantages of bMIND are more apparent in challenging but common settings, for instance, moderate effect size and more cell types. Interestingly, the CTS-DE analysis based on bMIND is robust to the specification of rough prior distributions (bMIND rp). As expected, the power of CTS-DE analysis increases with the effect size differentiating cases and controls (Fig. 3d) and with the number of samples evaluated (Fig. 3f), but decreases as the number of cell types estimated increases (Fig. 3e).

**Figure 3:**
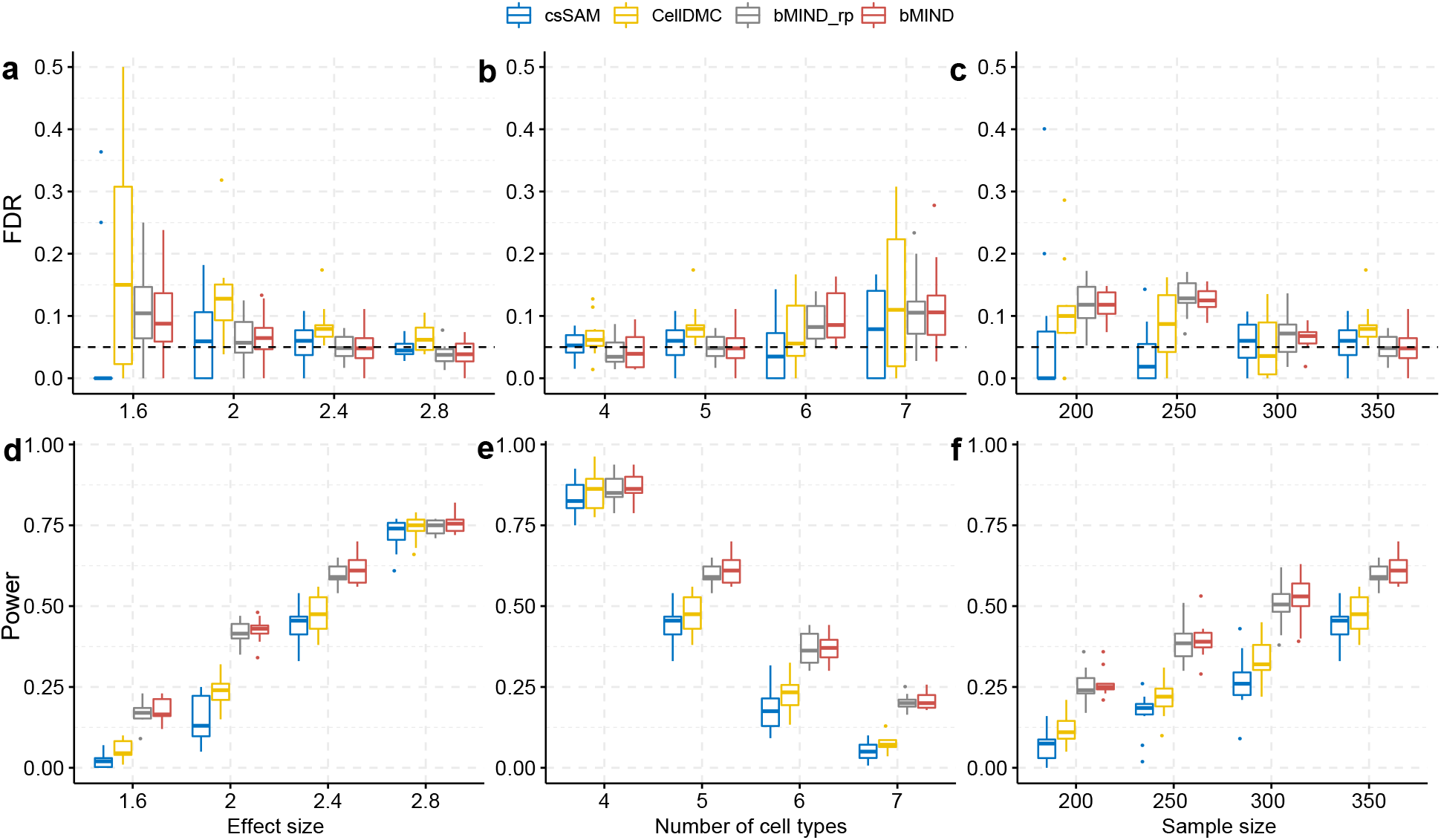
FDR and power comparison for CTS differential testing. **a-c**) FDR as a function of effect size in differentially expressed genes (**a**), the number of cell types (**b**), and the number of samples (**c**). **d-f**) Power as a function of effect size (**d**), the number of cell types (**e**), and the number of samples (**f**). All simulation scenarios are replicated ten times. If not specified, the effect size is set as 2.4, number of cell types *K* = 5, number of samples *N* = 350, number of measures *T* = 1, and 2% of gene-cell-type pairs with signal. The nominal FDR threshold is 0.05. bMIND rp: bMIND with a rough prior distribution.

### CTS differential expression analysis of ASD brain tissue

The snRNA-seq Velmeshev data provided an ideal resource for analyzing bulk RNA-seq data related to autism. With these data as reference, we analyzed the PsychENCODE UCLA-ASD bulk RNA-seq cortex data, also obtained from brain tissue samples from subjects diagnosed with ASD and from control subjects (Parikshak et al., 2016). First, we estimated cell type fractions using Bisque (Jew et al., 2020) and then inferred CTS using bMIND. Similar to the findings based on snRNA-seq data (Velmeshev et al., 2019), we found that there were more astrocytes and fewer oligodendrocytes in ASD than control samples (Fig. 4a). Because microglia and endothelial cells showed average fractions below 0.05 (Velmeshev et al., 2019), we dropped these cell types before differential expression analysis. Using bMIND we identified 688 CTS-DEGs over five major cell types (FDR < 0.05 and absolute log2 fold change > 0.14; Supplemental Table S1). For comparison, analysis of the 41 snRNA-seq samples produced 513 CTS-DEGs (Velmeshev et al., 2019) with the same criteria. Most of the CTS-DEGs identified by bMIND were from excitatory neurons (Fig. 4b), which concurs with the snRNA-seq findings (Velmeshev et al., 2019). In contrast, CellDMC (Zheng et al., 2018) identified 5,631 DEGs in inhibitory neurons and 2,502 DEGs in excitatory neurons (FDR < 0.05). However, CellDMC has been known to detect too many signals in analyses of real data (Rahmani et al., 2019), suggesting that the method is not robust to violations in the modeling assumptions.

**Figure 4:**
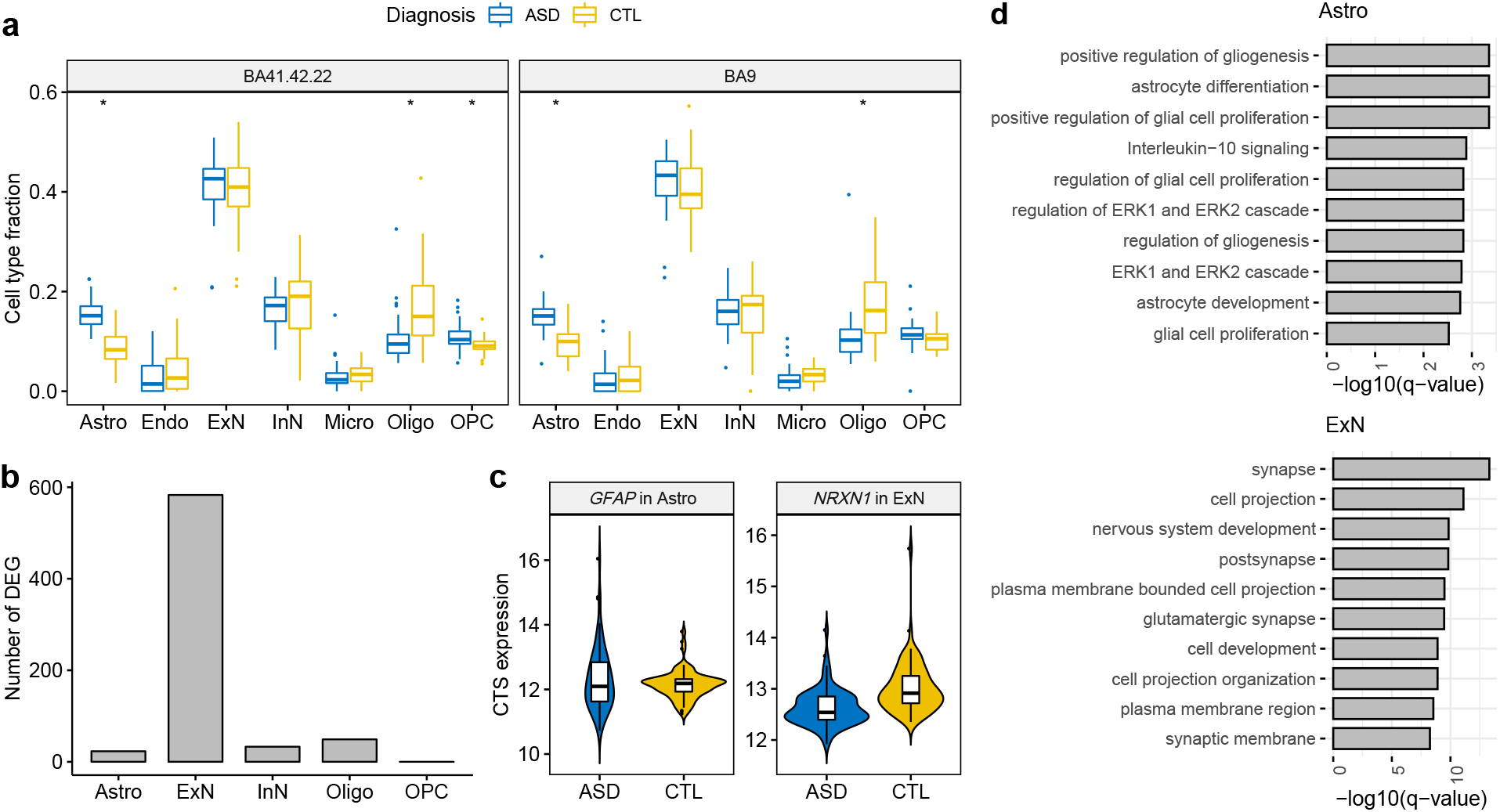
CTS differential expression analysis of autism. **a**) Estimated cell-type fractions for two cortical regions of the PsychENCODE UCLA-ASD data (Parikshak et al., 2016). * denotes significance after Bonferroni adjustment (p-value < 0.05/14) comparing fractions of ASD and control samples. **b**) Number of CTS-DEGs identified by bMIND in each cell type. **c**) Examples of bMIND identified CTS-DEGs. **d**) Gene ontology enrichment analysis for CTS-DEGs in astrocytes and excitatory neurons: top ten terms with FDR < 0.05.

When comparing CTS-DEGs detected by bMIND and snRNA-seq data (Velmeshev et al., 2019), and examining significant results found in excitatory neurons, we obtained 33 genes in common (Fisher’s exact test p-value = 4.3 × 10^−19^), including *NRXN1* (Fig. 4c). We also discovered some CTS-DEGs using bMIND that have not previously been identified by Velmeshev et al. (2019), for instance, astrocyte marker gene *GFAP* (Fig. 4c). Among the bMIND identified CTS-DEGs, six genes (*GFAP*, *NRXN1*, *LRRC4C*, *KCNMA1*, *RORB*, *SLC6A1*) were among the 102 ASD risk genes discovered by Satterstrom et al. (2020) (Fisher’s exact test p-value = 0.04) and 49 genes were among the SFARI autism gene list (Abrahams et al., 2013) (Fisher’s exact test p-value = × 10^−9^). As compared with the top 50 marker genes for each cell type derived from snRNA-seq data (Velmeshev et al., 2019), there was a significant enrichment of CTS-DEGs as markers in astrocytes and excitatory neurons (Fisher’s exact test p-value = 8.3 × 10^−4^ and 7.8 × 10^−8^, respectively).

We then evaluated the CTS-DEG sets with gene ontology (GO) enrichment analysis (Raudvere et al., 2019) (Fig. 4d and Supplemental Table S2). CTS-DEGs identified in astrocytes were significantly enriched in the regulation of gliogenesis, astrocyte differentiation/development, and glial cell proliferation. Correspondingly, the CTS-DEGs in excitatory neurons were enriched in glutamatergic (excitatory) synapse and nervous system development. We further parsed this set of enriched terms using REVIGO (Supek et al., 2011), a tool for clustering and interpreting long lists of GO terms. ASD DEGs were associated with 488 enriched GO terms. REVIGO identified two key themes for these terms, cell projection organization and neurotransmitter transport, as well as more minor themes of nervous system development and regulation of GTPase activity.

### CTS differential expression analysis of brain tissue from Alzheimer’s disease subjects

In the second case study, we conducted CTS-DE analysis for Alzheimer’s disease. We analyzed bulk RNA-seq data from brain samples of subjects diagnosed with AD and unaffected control subjects from three projects: Mayo Clinic RNA-seq data (Allen et al., 2016), MSBB (Mount Sinai Brain Bank) (Wang et al., 2018), and ROSMAP (Religious order Study and the Memory and Aging Project) (Bennett et al., 2018). We used AD snRNA-seq data (Mathys et al., 2019) for reference and the Bisque algorithm (Jew et al., 2020) to estimate cell type fractions. Following the cell clustering in the snRNA-seq data (Mathys et al., 2019), we focused on six cell types: Astro, ExN, InN, Micro, Oligo, and OPC.

Using bMIND, we estimated sample-level CTS expression and detected CTS-DEGs related to AD with FDR < 0.05 (Supplemental Table S1). Similar to the findings based on snRNA-seq of AD (Mathys et al., 2019), most identified CTS-DEGs were from excitatory neurons, a finding that comports with the observed selective vulnerability of excitatory neurons in the brain of AD samples (Leng et al., 2020). We compared the ExN-DEGs identified by the snRNA-seq study (Mathys et al., 2019) and bMIND from the three bulk datasets (Fig. 5a). The different numbers of DEGs can be explained by sample size and brain region heterogeneity. At the bulk expression level, an existing study (Coelho et al., 2020) also found more DEGs in the temporal lobe (Mayo data and MSBB Brodmann areas 22 and 36) than in frontal lobe (ROSMAP data and MSBB Brodmann areas 10 and 44). When we contrasted the ExN-DEGs found from snRNA-seq (Mathys et al., 2019), we observed significant overlap with bMIND ExN-DEGs for both the Mayo and MSBB data (Fisher’s exact test p-value = 3.9 × 10^−13^ and 1.8 × 10^−5^, respectively).

**Figure 5:**
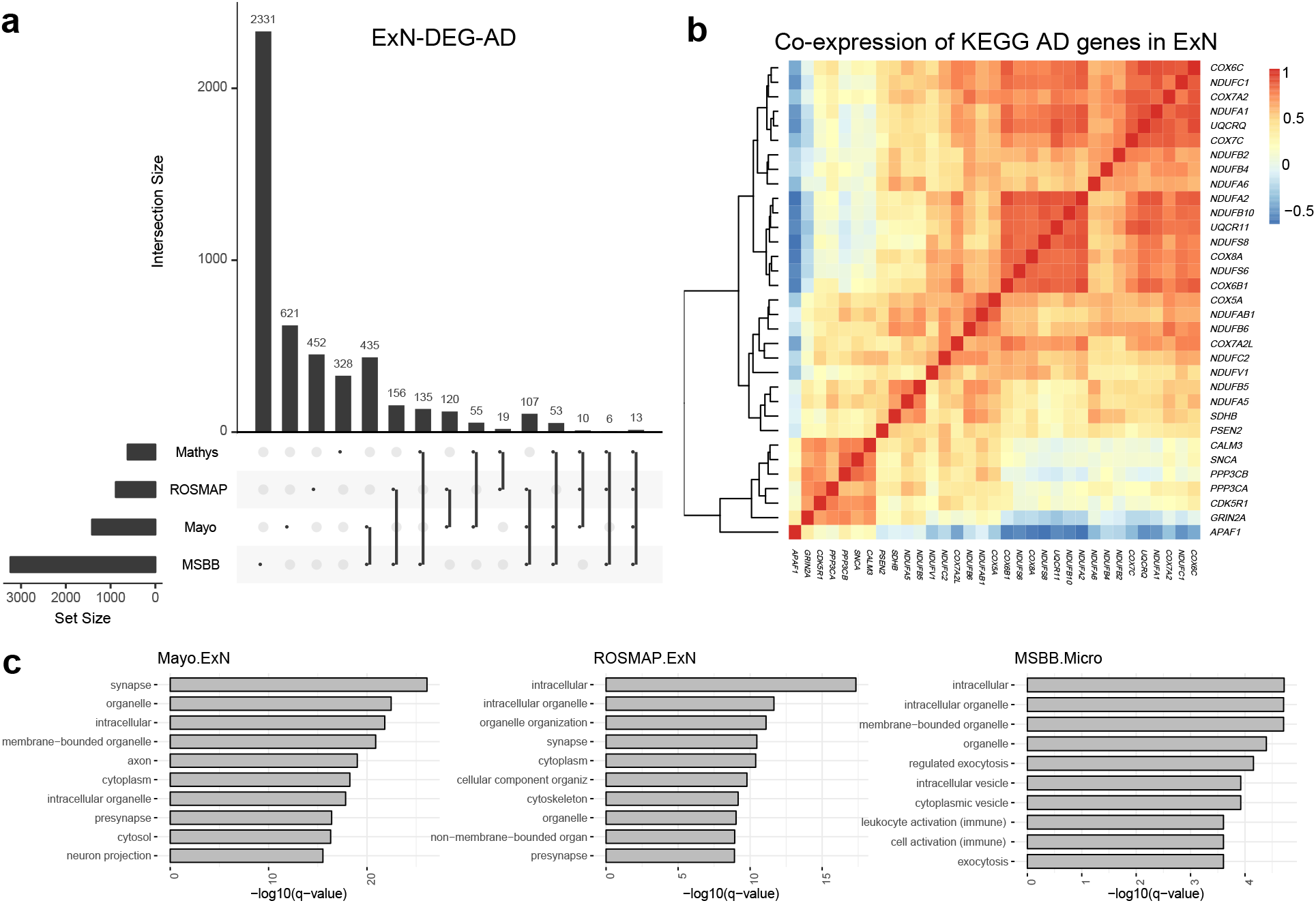
CTS differential expression analysis of Alzheimer’s disease. **a**) Intersection of DEGs in excitatory neurons (ExN-DEGs) related to AD identified in snRNA-seq data (Mathys et al., 2019) and three bulk RNA-seq datasets (ROSMAP, Mayo, and MSBB) by bMIND. **b**) Correlation matrix of the intersection of ExN-DEGs in Mayo data and the KEGG AD gene pathway. Correlation is computed using bMIND estimated sample-level CTS expression. **c**) Gene ontology enrichment analysis for ExN DEGs for Mayo and ROSMAP bulk data and microglia DEGs for MSBB data (MSBB.Micro). Here we present the top ten terms with FDR < 0.05 for each cell type.

Notably, CTS-DEGs in excitatory neurons identified by the Mayo data were enriched in KEGG Alzheimer’s disease pathway (Kanehisa and Goto, 2000) (Fisher’s exact test p-value = 1.5 × 10^−5^). To illustrate how genes worked together in Alzheimer’s disease at the cell type level, we took advantage of bMIND’s estimates of sample-level CTS expression to construct a co-expression network of a subset of genes expressed in excitatory neurons (Fig. 5b); here, the genes illustrated were those shared by the two gene sets, ExN-DEG-AD and KEGG Alzheimer’s disease pathway. We also conducted gene ontology enrichment analyses (Raudvere et al., 2019) for DEGs in different cell types (Supplemental Table S2). As expected, enriched terms for DEGs from excitatory neurons were enriched in synaptic and neuronal functions. For example, using REVIGO to assess the large number (683) of enriched GO terms for Mayo data identified several key themes: vesicle-mediated transport in the synapse, regulation of catabolism, and chemical synaptic transmission; and more minor themes of organelle organization and macro- and autophagy. DEGs in microglia, however, were enriched in immune processes (Fig. 5c).

### Contrasting functions of DEGs from excitatory neurons of ASD versus AD subjects

The bulk of DEGs for both ASD and AD arose from excitatory neurons, presenting an opportunity to learn more about both phenotypes based commonalities and differences of enriched GO terms. (Here we focused on DEGs from the temporal lobe of Mayo subjects.) ASD DEGs were enriched for 309 GO terms that were not shared with enriched terms for AD DEGs. From these terms, REVIGO identified one key theme, regulation of synapse organization, and two minor themes, amino acid transport and anatomical structure morphogenesis (Supplemental Fig. S2). This was somewhat different than the key themes associated with the entire set of ASD DEG, namely cell projection organization and neurotransmitter transport. In fact, all of these themes are likely important to liability for ASD (Satterstrom et al., 2020).

There were 504 specific enriched GO terms for AD DEGs, from which REVIGO identified key themes of vesicle-mediated transport and regulation of catabolism. More minor themes were organelle organization, macro- and autophagy, and protein/macromolecule modification (Supplemental Fig. S2). These themes were quite similar to those identified from the entire list of enriched GO terms for AD DEGs

Next, we asked how REVIGO interpreted the enriched terms that were shared between ASD and AD. Notably, while 179 GO terms were identical, the FDR q-values REVIGO used to prioritize them were not. To the contrary, there was no relationship between ASD and AD q-values for these shared terms (p-value = 0.68, paired Wilcoxon test). The major themes that emerged for ASD were cell projection organization and neurotransmitter transport, quite similar to those for the entire set of enriched GO terms associated with ASD DEGs. For AD, however, the major theme was regulation of cellular component organization, a theme that recurred across the different partitions of enriched GO terms for AD DEGs.

### CTS-eQTL analysis of GTEx V8 brain data

To generate a resource of inferred CTS-eQTLs for various brain regions, we analyzed the latest GTEx brain data (Aguet et al., 2019) (V8) using bMIND. We first obtained the cell type fractions for each GTEx bulk sample via non-negative least squares and signature matrix derived from Darmanis et al. (2015) and described in Wang et al. (2020). We then estimated subject-level CTS expression for 11 GTEx brain regions, after combining replicates for frontal cortex and cerebellum: amygdala, cerebellum, anterior cingulate cortex, frontal cortex, hippocampus, hypothalamus, substantia nigra, caudate, nucleus accumbens, putamen, and spinal cord. For each region, gene expression was estimated for six cell types: Astro, Endo, ExN, InN, Micro, and Oligo. Then, with genotypes and estimated subject-level region-specific CTS expression, we calculated cis-eQTLs for each brain region cell type using MatrixEQTL (Shabalin, 2012) for all variants with minor allele frequency > 1%. Cis was defined as +/− 1Mb around each gene. The summary statistics of significant (FDR < 0.05) gene-variant pairs were saved in GitHub folder: https://github.com/randel/bMIND_GTEx8-signif_region_CTS_eQTLs_cis.

To evaluate the results of eQTL mapping, we first confirmed that the eQTL analysis p-values were well-calibrated (Fig. 6a, Supplemental Fig. S3) and eQTLs were enriched near the transcriptional start site (TSS), as expected (Fig. 6b, Supplemental Fig. S4). Next, we hypothesized that many of our region-specific CTS eQTLs would match the GTEx regional analysis of eQTLs using bulk data (Aguet et al., 2019). To make this comparison, we calculated the fraction of bulk eQTLs per region as detected as region-specific CTS eQTLs by bMIND (Fig. 6c), noting substantial concordance in general. In addition, as might be expected, the eQTL mapping fractions were highly correlated with the average cell type fraction per region, with a Pearson correlation of 0.88. The high concordance reveals the important role of cell type abundance in bulk data analysis, and both analyses verify the replicability of our CTS analysis.

**Figure 6:**
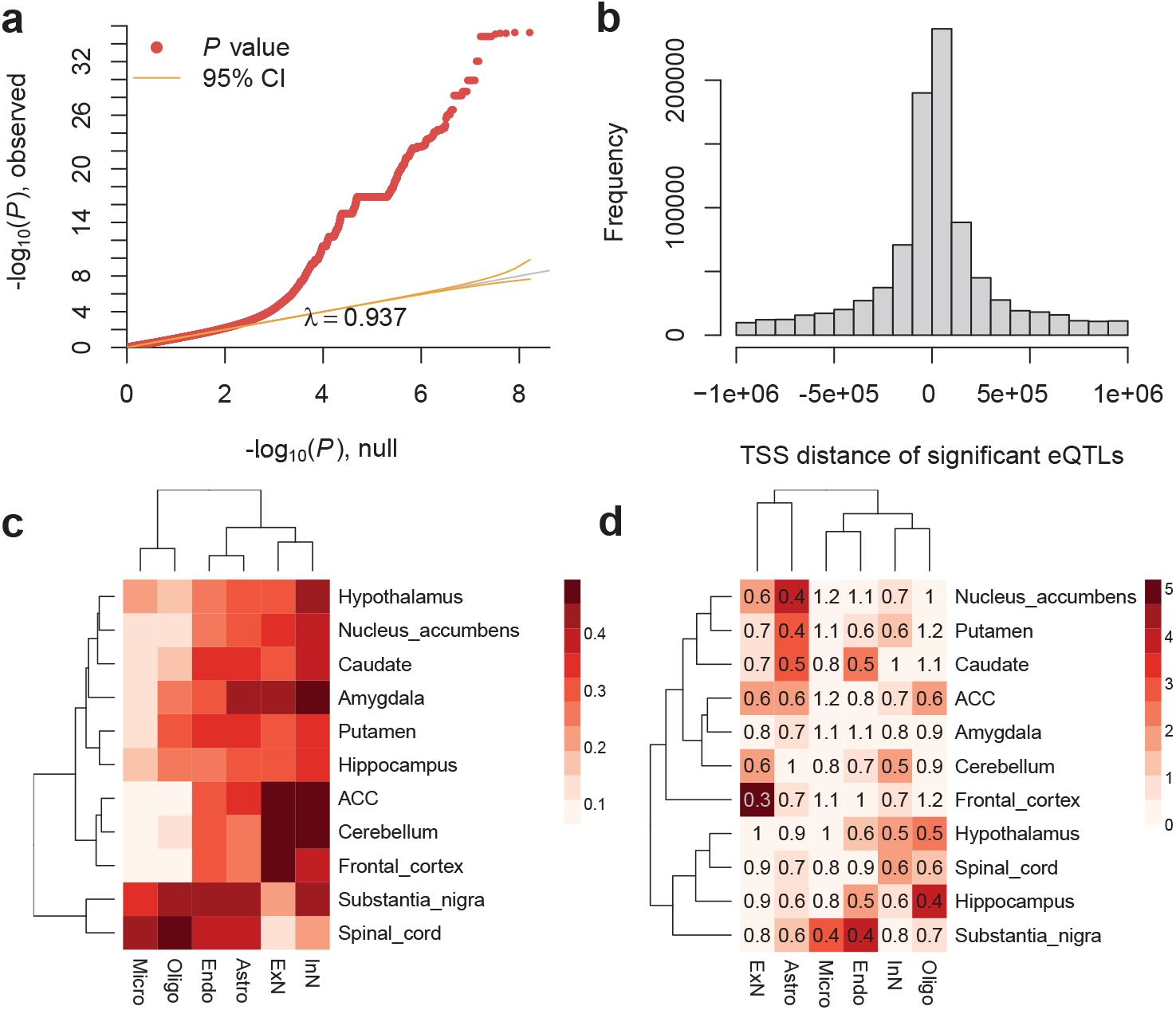
Region-specific CTS eQTL analysis with GTEx V8 brain data. **a)** The QQ-plot of p-values from eQTL analysis. Here we show an example of microglia in substantia nigra. **b)** The enrichment of significant eQTLs near TSS. Here we present an example of excitatory neurons in the frontal cortex. **c**) Fraction of GTEx brain bulk eQTLs detected as CTS eQTLs in each brain region. ACC: anterior cingulate cortex. **d)** The enrichment analysis of ASD genes in region-specific CTS eGenes (genes with eQTLs). The heatmap color denotes −log10 transformed Benjamini-Hochberg adjusted p-values (Benjamini and Hochberg, 1995) based on two-sided Fisher’s exact tests, and the number represents the odds ratio.

To assess the utility of bMIND’s region-specific CTS eQTLs, we assessed the connection between ASD genes (Satterstrom et al., 2020) and genes with eQTLs (eGenes). Using pLI score (Lek et al., 2016) as a measure of gene conservation, we first replicated a previous finding that eQTLs do not tend to occur in very conserved genes (Lek et al., 2016; Werling et al., 2020). For instance, in ExN of frontal cortex, the odds ratio of being eGenes and conserved (pLI ≥ 0.995) was 0.53 (Fisher’s exact test p-value = 1.2 × 10^−31^). Given that conserved genes were less likely to be eGenes and ASD genes tended to be very conserved (Satterstrom et al., 2020), it is reasonable to predict that ASD genes were less likely to be eGenes. This speculation was verified with enrichment analysis of ASD genes as eGenes. Curiously, relatively fewer ASD genes (Satterstrom et al., 2020) were likely to be eGenes in excitatory neurons of the frontal cortex (Fig. 6d), which thus far is the most related cell type and brain region associated with autism. In large part, this pattern emerges because ASD genes tend to have higher expression levels, as do cortical excitatory neurons, and ASD genes tend to be highly conserved.

## Discussion

We develop a Bayesian algorithm, bMIND, to provide CTS expression for bulk RNA-seq samples with prior information derived from scRNA-seq data. This approach addresses the limitation of bulk RNA-seq, namely it arises as a convolution of cell-type gene expression profiles, and the limitation of scRNA-seq data, namely ample technical noise and a limited number of samples. Yet, bMIND builds on the unique advantages of each data type, which include a large sample size of bulk RNA-seq and cell-type gene expression profiles from scRNA-seq. We conduct extensive simulations to compare bMIND with state-of-the-art methods, demonstrating that bMIND improves the estimation accuracy and differential testing power, while controlling FDR. Through analysis of CTS differential expression for brain samples of subjects diagnosed with ASD or unaffected, as well as a similar design for Alzheimer’s disease, we show the utility of bMIND to enhance the understanding of etiology with cell-type resolution. Finally, by analysis of the latest GTEx V8 data using bMIND, we obtain CTS eQTLs for eleven brain regions. To the best of our knowledge, this is the most comprehensive brain CTS eQTL resource, with which we verify existing findings and believe will prove valuable for numerous studies.

Nonetheless, bMIND inevitably has limitations. For instance, bMIND provides more accurate CTS estimates for more abundant cell types, and thus in those cell types the CTS differential expression analysis should be more powerful, all other things being equal. This issue also applies to the CTS differential expression analysis using cell-level data from scRNA/snRNA-seq, where cell types with more cells/nuclei quantified have much more power to detect CTS-DEGs. More work will be needed to develop methods sensitive to less common cell types. In future work, we also plan a fully Bayesian approach to account for all sources of variation, such as the variation involving estimation of cell type fractions. Similar to other CTS analysis methods (Luo et al., 2020), bMIND is robust to moderate estimation error in cell type fractions. Here we focus on RNA-seq data, but the approach for bMIND could also be used for the analysis of other omics data, such as DNA methylation. We will pursue this direction in future work. When there are no scRNA-seq data available, we can use those parameter estimates from MIND (Wang et al., 2020) as hyperparameters in the prior distribution.

Note that CTS differential expression analysis is different from cell type enrichment analysis (Skene and Grant, 2016). CTS differential expression analysis not only links diseases to specific cell types, but it also deepens the analysis to identify certain disease-related genes within those cell types. While it remains a challenging task, we show the advantage and flexibility of estimating the virtual CTS expression profile for each bulk RNA-seq sample. In addition to the improved power to detect CTS DEGs and eQTLs, sample-level CTS expression enables the development of co-expression networks specific to certain cell types and other sample-level analyses.

## Materials and methods

### Software implementation

We implement the Bayesian mixed-effects models in bMIND with the MCMCglmm algorithm (Hadfield, 2010), which fits a broad class of Bayesian generalized linear mixed-effects models based on MCMC approaches. It is flexible to incorporate normal prior for fixed effects and inverse-Wishart prior for the covariance matrix. As the conjugate prior, the inverse-Wishart prior facilitates estimation and allows the incorporation of the prior cell-type covariance matrix estimated from scRNA-seq data explicitly. To make the computation feasible for all genes in the genome, we analyze one gene each time and run the analysis in parallel. To build a user-friendly software package, we integrate the following two steps: estimating cell-type fraction and CTS expression. With users’ input of bulk data and either raw scRNA-seq data reference or signature matrix, we can output both cell type fractions via non-negative least squares or Bisque (Jew et al., 2020) and CTS expression via bMIND. When a phenotype is provided, the package will conduct CTS-DE analysis and output p-values adjusted for multiple testing. The R package is publicly hosted on GitHub (https://github.com/randel/MIND) with detailed tutorials.

### CTS differential expression analysis

The output of bMIND is a 3-dimensional array (gene × cell type × sample). With estimated sample-level CTS expression, we are able to conduct analyses that are previously only available using bulk RNA-seq data, deepening the analyses from tissue level to cell-type level. Here we focus on CTS differential expression (DE) analysis as an example. To control for false discovery rate (FDR) in CTS-DE analysis, we propose a stringent testing procedure:

1. We first conduct the multivariate analysis of variance (MANOVA) using CTS expression for each gene with respect to the phenotype of interest and claim a gene as a DEG by Benjamini-Hochberg adjusted p-values (Benjamini and Hochberg, 1995).
2. To find in which cell type a DEG is differentially expressed, we obtain CTS-DE p-values by regressing the phenotype on CTS expression in that gene.
3. A CTS-DEG is determined if the CTS-DE p-value is the minimal across cell types in a DEG and less than 0.05/*K*, where *K* is the number of cell types (Guo et al., 2010).

That is, we only detect the top significant signal across cell types within each gene. Existing snRNA-seq studies of ASD and Alzheimer’s disease (Mathys et al., 2019; Velmeshev et al., 2019) support this testing scheme that most (79-93%) CTS-DEGs are only differentially expressed in a single cell type.

In the simulation study, we assess the FDR and power as a function of effect size, the number of cell types, and the number of samples. All simulation scenarios are replicated ten times. For each replicate, we simulate 500 genes expressed in *K* cell types (*K* = 5 except for the scenario with varying *K*) for 350 samples (except when we vary the sample size). We randomly select 2% of the 500 × *K* gene-cell-type combination as differentially expressed genes in a cell type, with an effect size of 2.4 (mean difference divided by the standard deviation of bulk expression), except when the effect size is varied. The data generation model is adapted from the csSAM R package (Shen-Orr et al., 2010), but we add sample-level variability for CTS expression.

### Data resources and analyses

We utilized public bulk and single-nucleus RNA-seq (snRNA-seq) datasets. For analysis of ASD, we used bulk RNA-seq data from the PsychENCODE UCLA-ASD project (Parikshak et al., 2016), which collected 167 tissue samples from 2 cortical regions of 91 subjects (47 ASD and 44 control subjects). Those subjects ranged in age at death from 2 to 67, with 22 being the median age. Most (81%) of those subjects were male. As a reference of cell type information, we used snRNA-seq data from an ASD study (Velmeshev et al., 2019), which collected 105 thousand nuclei from 41 cortical samples taken from both ASD and control subjects. We also used the snRNA-seq data (Velmeshev et al., 2019) to generate realistic bulk data for the simulation studies. To evaluate the consistency of estimation results, we utilized two similar bulk RNA-seq datasets (Parikshak et al., 2016; Velmeshev et al., 2019).

For the case study of Alzheimer’s disease, we used bulk RNA-seq data from three resources: the Mayo Clinic (Allen et al., 2016) data with 160 samples from the temporal cortex, MSBB (Mount Sinai Brain Bank) (Wang et al., 2018) data with 850 bulk samples from 4 cortical regions (Brodmann areas 10, 22, 36, and 44), and ROSMAP (Religious Order Study and the Memory and Aging Project) (Bennett et al., 2018) with 636 samples from the dorsolateral prefrontal cortex. We adopted a consistent definition for Alzheimer’s disease (Braak score ≥ 4) across datasets. We compared our CTS-DEGs with those from the snRNA-seq study (Mathys et al., 2019), which quantified the expression of 81 thousand nuclei from the cortex of 48 subjects. All bulk RNA-seq and snRNA-seq data include both affected and control subjects.

Sets of DEG from both ASD and AD were analyzed for functional effects as determined by gene ontology (GO) enrichment analysis (Raudvere et al., 2019), using threshold FDR *q* < 0.05. To capture the major functions of the DEGs obtained from ASD and from the Mayo study of AD, we analyzed their enriched GO terms by REVIGO (Supek et al., 2011). REVIGO assesses semantic similarity of GO terms (Schlicker et al., 2006), clusters similar terms, prioritizes more enriched terms for the semantic interpretation, and displays representative terms for the cluster. For these analyses, we used a similarity setting of 0.5, which favors shorter and semantically diverse lists of functions, as well as two default settings: semantic similarity measure SimRel and the database for GO term sizes “whole UniProt”. The terms were analyzed by the online version of REVIGO, which used GO release “go monthly-termdb.obo-xml.gz” (Jan 2017) and UniProt-to-GO mapping file “goa uniprot gcrp.gaf.gz” (15 Mar 2017).

## Acknowledgements

The results published here are in part based on data obtained from the AMP-AD Knowledge Portal. Mayo RNA-seq data were provided by the following sources: The Mayo Clinic Alzheimers Disease Genetic Studies, led by Dr. Nilufer Taner and Dr. Steven G. Younkin, Mayo Clinic, Jacksonville, FL using samples from the Mayo Clinic Study of Aging, the Mayo Clinic Alzheimers Disease Research Center, and the Mayo Clinic Brain Bank. Data collection was supported through funding by NIA grants P50 AG016574, R01 AG032990, U01 AG046139, R01 AG018023, U01 AG006576, U01 AG006786, R01 AG025711, R01 AG017216, R01 AG003949, NINDS grant R01 NS080820, CurePSP Foundation, and support from Mayo Foundation. Study data includes samples collected through the Sun Health Research Institute Brain and Body Donation Program of Sun City, Arizona. The Brain and Body Donation Program is supported by the National Institute of Neurological Disorders and Stroke (U24 NS072026 National Brain and Tissue Resource for Parkinsons Disease and Related Disorders), the National Institute on Aging (P30 AG19610 Arizona Alzheimers Disease Core Center), the Arizona Department of Health Services (contract 211002, Arizona Alzheimers Research Center), the Arizona Biomedical Research Commission (contracts 4001, 0011, 05-901 and 1001 to the Arizona Parkinson’s Disease Consortium) and the Michael J. Fox Foundation for Parkinsons Research. MSBB data were generated from postmortem brain tissue collected through the Mount Sinai VA Medical Center Brain Bank and were provided by Dr. Eric Schadt from Mount Sinai School of Medicine. ROSMAP data were provided by the Rush Alzheimer’s Disease Center, Rush University Medical Center, Chicago. Data collection was supported through funding by NIA grants P30AG10161 (ROS), R01AG15819 (ROSMAP; genomics and RNAseq), R01AG17917 (MAP), R01AG30146, R01AG36042 (5hC methylation, ATACseq), RC2AG036547 (H3K9Ac), R01AG36836 (RNAseq), R01AG48015 (monocyte RNAseq) RF1AG57473 (single nucleus RNAseq), U01AG32984 (genomic and whole exome sequencing), U01AG46152 (ROSMAP AMP-AD, targeted proteomics), U01AG46161(TMT proteomics), U01AG61356 (whole genome sequencing, targeted proteomics, ROSMAP AMP-AD), the Illinois Department of Public Health (ROSMAP), and the Translational Genomics Research Institute (genomic). Additional phenotypic data can be requested at www.radc.rush.edu.

The Genotype-Tissue Expression (GTEx) Project was supported by the Common Fund of the Office of the Director of the National Institutes of Health (commonfund.nih.gov/GTEx). Additional funds were provided by the NCI, NHGRI, NHLBI, NIDA, NIMH, and NINDS. Donors were enrolled at Biospecimen Source Sites funded by NCI\Leidos Biomedical Research, Inc. subcontracts to the National Disease Research Interchange (10XS170), Roswell Park Cancer Institute (10XS171), and Science Care, Inc. (X10S172). The Laboratory, Data Analysis, and Coordinating Center (LDACC) was funded through a contract (HHSN268201000029C) to the The Broad Institute, Inc. Biorepository operations were funded through a Leidos Biomedical Research, Inc. subcontract to Van Andel Research Institute (10ST1035). Additional data repository and project management were provided by Leidos Biomedical Research, Inc.(HHSN261200800001E). The Brain Bank was supported supplements to University of Miami grant DA006227. Statistical Methods development grants were made to the University of Geneva (MH090941 & MH101814), the University of Chicago (MH090951, MH090937, MH101825, & MH101820), the University of North Carolina - Chapel Hill (MH090936), North Carolina State University (MH101819), Harvard University (MH090948), Stanford University (MH101782), Washington University (MH101810), and to the University of Pennsylvania (MH101822). The datasets used for the analyses described in this manuscript were obtained from dbGaP at http://www.ncbi.nlm.nih.gov/gap_through_dbGaP_accession_number_phs000424.v8.p2. This work was supported, in part, by National Institute of Mental Health (NIMH) grants R01MH123184, R37MH057881, R37MH057881-22S1, and by Simons Foundation Autism Research Initiative (SFARI) grant SF575547.

